# Transcranial direct current stimulation elevates the baseline activity while sharpening the spatial tuning of the human visual cortex

**DOI:** 10.1101/2023.02.07.527461

**Authors:** Jeongyeol Ahn, Juhyoung Ryu, Sangjun Lee, Chany Lee, Chang-Hwan Im, Sang-Hun Lee

## Abstract

**Background:** Although transcranial direct current stimulation (tDCS) is widely used to affect various kinds of human cognition, behavioral studies on humans have produced highly inconsistent results. This requires a clear understanding of how tDCS impacts the system-level neural activity, a prerequisite for the principled application of tDCS to human cognition.

**Objective:** Here, we aim to gain such understanding by probing the spatial and temporal cortical activity of the human early visual cortex (EVC) in diverse aspects while controlling the polarity and presence of tDCS. We target EVC to capitalize on its well-established anatomical and functional architecture that is readily accessible with non-invasive quantitative neuroimaging methods.

**Methods:** To create an electric field in EVC precisely and effectively, we tailored high-definition stimulation montages for 15 individual brains by running electric field simulations. We then conducted an fMRI (functional magnetic neuroimaging)-tDCS experiment on each brain with a sham-controlled crossover design over multiple days. We quantified tDCS effects with eight measures, tested their significance with mixed ANOVA, and further validated their robustness to across-voxel and across-subject variability.

**Results:** The anodal application of tDCS gradually elevated EVC’s baseline BOLD activity and sharpened its spatial tuning by augmenting surround suppression without affecting its evoked activity.

**Conclusions:** Comparisons of our and previous findings suggest the fundamental differences in tDCS effects between the visual and motor cortices, inhibitory and excitatory effects predominant in the former and latter, respectively. This calls for considering the differences in the excitatory-inhibitory recurrent network between brain regions in predicting or interpreting tDCS effects.

## Introduction

Early studies on animal brains [1,2] suggested that surface-positive and negative currents elevate and suppress neuronal excitability. These effects on neural excitability lasted for several minutes to hours after the stimulation was switched off. However, the effects of transcranial direct current stimulation (tDCS) on human behavior are inconsistent. The polarity-specific modulation of excitability seems symmetric in the motor-domain behavior but asymmetric in other cognitive-domain behaviors: upregulation by anodal-tDCS (a-tDCS) was often significant, whereas downregulation by cathodal-tDCS (c-tDCS) was rarely significant [3]. Furthermore, a recent meta-analysis of tDCS studies indicates that even a-tDCS shows inconsistent results across studies [4].

Many sources could have contributed to the inconsistency across human studies on tDCS. First, unlike TMS or DBS, tDCS does not induce massive neural activities but modulates spontaneous or evoked neural responses [5–8]. Thus, tDCS effects depend on whether a target cortical region is in an active state. For example, the tDCS effects on working memory depended on task difficulty [9] and the idiosyncratic level of baseline performance [9,10]. Likewise, the behavioral effects of the tDCS on the motor cortex depended on task types [11–13]. Second, the inconsistency might arise from individual differences at various levels, including brain and skull morphology, neurotransmitter composition, and genetic profile [14,15]. In support of this, the electric-field differences owing to skull and brain anatomies [16,17] or neurotransmitter efficiency [18] accounted for the across-individual variability of tDCS effects on behavioral or neural measurements. Lastly, poor experimental designs are also a source of inconsistency. Any failure in controlling the various within-day and across-day noises and in making both subjects and experimenters blind to stimulation conditions is likely to mask subtle, modulatory tDCS effects [19,20].

Given these various interfering sources, the inconsistency in tDCS effects on human cognition requires a clear understanding of how tDCS perturbs the cortical activities of the brain regions underlying any given performance [21]. Without this understanding, whatever behavioral effects of tDCS are almost always subject to the interfering sources that interact with the brain activity, which it difficult to interpret any observed behavioral effects. To gain such understanding, we acquired blood-oxygenation-level-dependent (BOLD) responses to dynamic visual patterns from the human early visual cortex (EVC) using functional magnetic resonance imaging (fMRI) while stimulating the same cortex with tDCS.

To effectively examine the tDCS effects on brain activity while addressing the interfering sources, we incorporated the following points into the design of our experiment. First, we targeted EVC because its well-known functional architecture, such as retinotopic representations by neurons with receptive fields [22–25], allows us to estimate various system-level changes of cortical activity. Such a refined assessment of tDCS effects would provide informative clues for how tDCS affects cortical activity. Second, to create an ideal electric field in EVC, we used a multi-electrode tDCS system in conjunction with a high-fidelity stimulation protocol. Specifically, to address the across-subject differences due to skull and brain morphology, we tuned stimulation protocols using the electric field simulation carried out on the individual head anatomies. Lastly, we adopted a sham-controlled crossover design, such that each subject participated in all three types of the daily session—the ‘anodal,’ ‘cathodal,’ and ‘sham’ sessions. All sessions shared an identical structure, acquiring BOLD responses before (pre-stimulation), during (peri-stimulation), and after (post-stimulation) stimulation. This design corrects the BOLD responses in the four main experimental conditions (2 stimulation polarities x 2 stimulation phases) for the tDCS-irrelevant fluctuations in BOLD signal known to occur within a daily session and between daily sessions or individuals.

We assessed the tDCS effects on EVC for the following aspects. We inspected the temporal dynamics of the BOLD responses to a transient full-field visual pattern to see whether tDCS perturbs the transient or sustained neural activity or the neuro-vascular function. We also inspected the spatial tuning of EVC using traveling-wave visual patterns [26–30]. Changes in pRF would inform us of the cortical mechanism that mediates tDCS effects, such as the surround inhibition or long-range facilitation mechanisms known to modulate the spatial tuning of visual neurons [31–34]. We also analyzed the noise correlation between cortical sites, which is known to play crucial roles in information processing as demonstrated by computational [35], animal electrophysiological [36], and human fMRI studies [37–39].

We found that a-tDCS elevated the baseline, but not evoked, cortical activity while sharpening the spatial tuning by augmenting surround suppression in EVC. These findings are at odds with the previous findings reported on the motor cortex, suggesting that tDCS effects substantially differ between the visual and motor cortices.

## Methods

### Subjects

Fifteen subjects (two authors included) with a mean age of 25.7 ± 4.17 years (five females) contributed to the dataset for analysis (see Supplementary materials for details). All subjects had normal or corrected-to-normal vision. They all provided written informed consent. All procedures complied with the safety guidelines and standards, as approved by the Institutional Review Board of Seoul National University (1711/003-027).

### fMRI data preprocessing

The images acquired across different sessions and runs were aligned to the high-resolution T1 images based on the T images (see Supplementary materials for details) using SPM8 (http://fil.ion.ucl.ac.uk/spm) and mrTools (http://cns.nyu.edu/heegerlab/?page=software). The first-cycle data of each run were discarded for fMRI signal stabilization, so the images acquired during the brief stimulation period in the sham scan runs were not included in the data analysis (Fig 1.d).

**Fig. 1.**
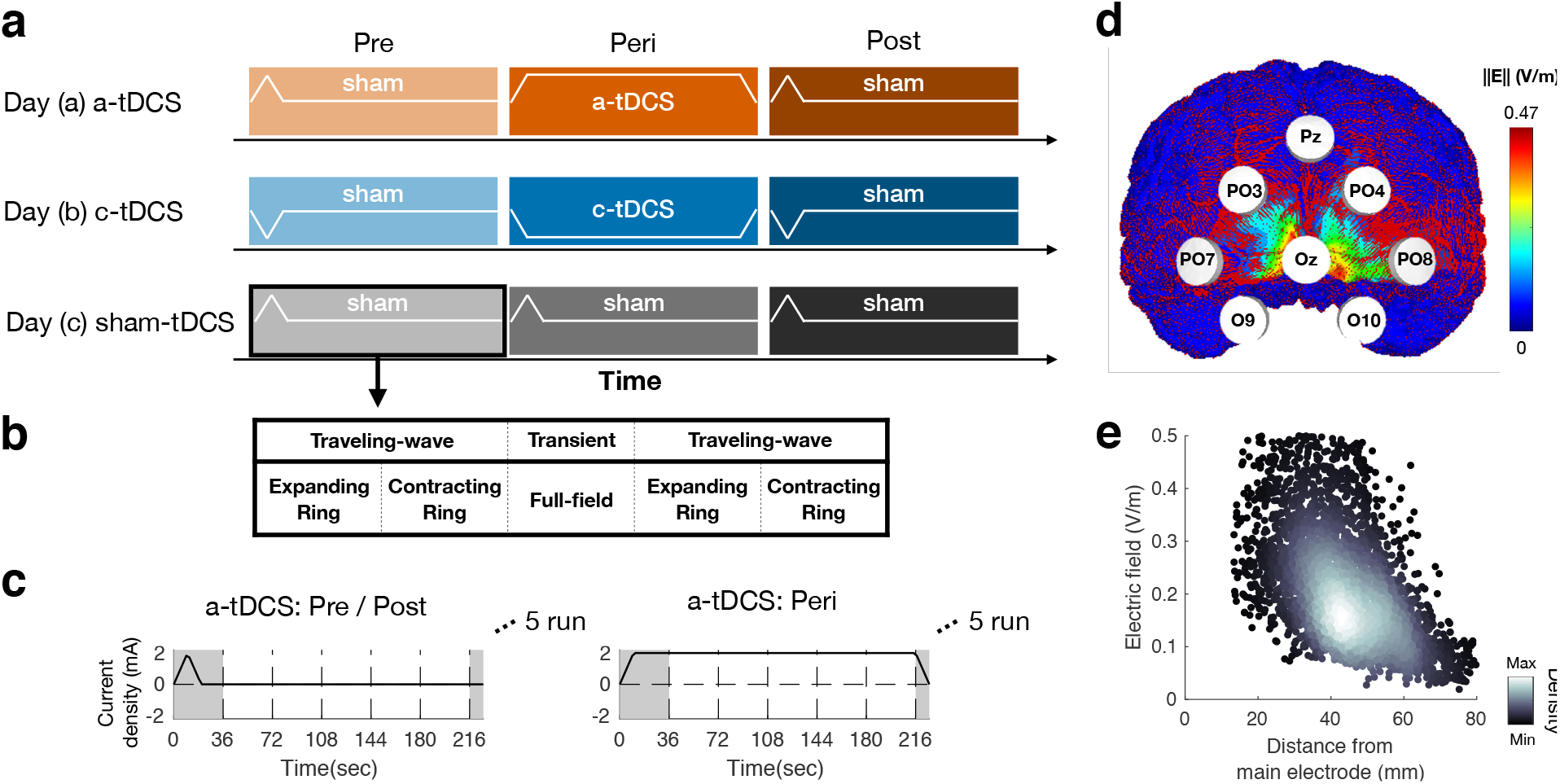
Design of tDCS-fMRI experiment and tDCS montage (a) Sham-controlled crossover design. Each subject participated in the anodal (a-tDCS), cathodal (c-tDCS), and sham (sham-tDCS) tDCS sessions on three different days. Each session consisted of three stimulation phases, one before (Pre), one during (Peri), and the other after (Post) the stimulation. (b) Structure of scan runs. Each stimulation phase consisted of five scan runs, four devoted to pRF mapping based on the traveling-wave input and the other to temporal profile mapping based on the transient full-field input. (c) Time courses of electric current. Two general types of electric stimulation were used (see the white line curves in a): the one used in Pre- or Post-stimulation phases and the other in Peri-stimulation phases. In the former, the current was switched on and quickly off as a control for placebo effects. In the latter, the current was switched on and maintained throughout the entire run. We discarded the BOLD data acquired during the first cycle in which the “fake-stimulation” was applied (demarcated by gray regions). (d) Eight-channel tDCS montage and an example electric field simulation shown for one subject. (e) Density plot of electric field density distribution estimated from simulation for all voxels.

### Experimental design: sham-controlled crossover experimental design with double blinds

Each subject participated in four daily sessions: retinotopy-mapping, a-tDCS, c-tDCS, and sham-tDCS sessions, which were randomized in order across subjects and separated from one another by at least a week to ensure a sufficient washout of the preceding effects. In each session a pre-stimulation phase was followed by peri-stimulation and post-stimulation phases, which are all identical in fMRI scan structure, each consisting of one temporal-profile scan and four spatial-profile scans (Figure 1.b). The genuine tDCS was applied only in the ‘peri-stimulation’ phase of the a-tDCS and c-tDCS sessions.

### tDCS setup

See the supplementary materials for how we optimized tDCS protocols for individual subjects using electrical field simulation and how we predicted the electric fields that are instigated by those tDCS protocols.

### Temporal-profile analysis of BOLD responses to transient full-field input

In each 36-second cycle, sinusoidal radial gratings were presented within a large (8° in radius) circular aperture for 3 seconds and then disappeared for 33 seconds (Fig 2.a.). We fit the hemodynamic impulse response function (HIRF) with the two gamma functions [44] to each voxel’s time series of BOLD responses by predicting the time series based on the convolution of the stimulus event matrix with the HIRF. From the best-fit predicted time course of BOLD responses, we defined ‘baseline’ as the predicted BOLD response at stimulus onset; ‘peak amplitude’ as the fractional change from the BOLD value at stimulus onset to that at peak (i.e., (*BOLD_peak_* − *BOLD_onset_*)/ *BOLD_onset_*); ‘Time to peak’ as the time taken for BOLD responses to reach the peak; ‘FWHM’ as the time taken to reach its second half-of-maximum value after it reached its first half-of-maximum value (Fig. 2.c).

**Fig. 2.**
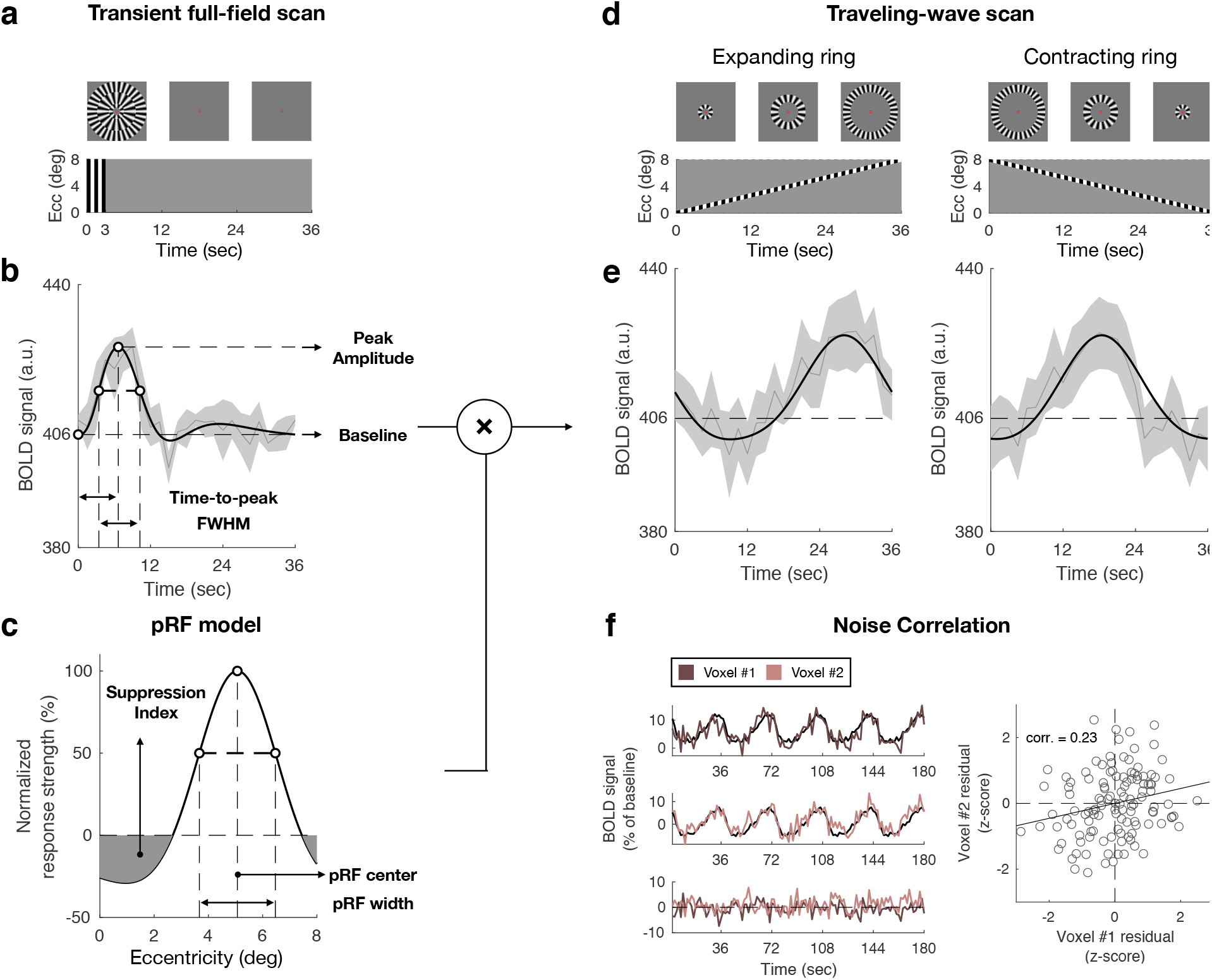
Quantification of temporal and spatial profiles of BOLD responses (a-b) Stimuli and paradigm for characterizing the spatial and temporal profile of BOLD responses to transient full-field (a) and traveling-wave (b) input. (c-d) Example one-cycle averages of a single voxel’s BOLD responses to the transient full-field (c) and traveling-wave (b) input. The model time series (black curves) was fit to the observed ones (gray curves). The shades demarcate the across-cycle standard deviation. The dashed lines and arrows indicate the temporal-profile measures in c. (e) Example difference-of-Gaussian model of pRF with three parameters fit to the single voxel’s BOLD responses to the traveling-wave input depicted in d. The dashed lines and arrows indicate the pRF measures. (f) Quantification of noise correlation. Left, the process of extracting noise time series shown for two example voxels. Noise time series were defined by subtracting the concatenated across-cycle averages of BOLD responses from the raw BOLD time series. Right, definition of noise correlation. The noise time points of one of the two voxels depicted in the left panel are plotted against those of the other voxel.

### Spatial-profile analysis of BOLD responses to traveling-wave input

In each 36-second cycle, sinusoidal radial gratings presented within ring-shape apertures slowly (4.5 v.a.d./sec) traversed the visual field in the radial direction (Fig. 2.b.). With the 1D difference-of-Gaussian (DoG) function [45], pRF was defined as a gain function (*g*(*x*)) of stimulus input (*x*) as follows:

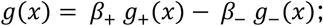

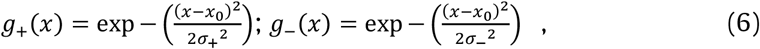

where *g*_+_(*x*) and *g*_−_(*x*) are positive and negative Gaussian functions, the respective contributions of which to *g*(*x*) are expressed by the two coefficients, *β*_+_ and *β*_−_; *x*_0_ is the center of pRF, and *σ*_+_ and *σ*_−_ are the standard deviations of the positive and negative Gaussian functions, respectively. Next, we predicted the aggregated responses of a population of visual neurons within a voxel to the traveling-wave input (*r*(*t*)) by convolving a binary matrix of spatiotemporal stimulus input (*s*(*x*, *t*)) with *g*(*x*) (Fig 2.c-e.). Finally, we predicted the BOLD responses by convolving *r*(*t*) with the HIRF defined by the transient-stimulus scan in each stimulation phase. From the best-fit pRF defined above, we defined ‘pRF center’ as the pRF parameter *x*_0_; ‘pRF width’ as the FWHM of the positive part of pRF; ‘pRF SI’ as the ratio of the integration of the negative part of pRF to the integration of the positive part of pRF within the visual field of interest (0-8 v.a.d. in eccentricity) (Fig. 2.e).

### Noise correlation estimation

We extracted the noise time series for each voxel by repeatedly concatenating the across-cycle average response to match the scan run length and subtracting it from the original BOLD. Then, Pearson’s correlation coefficient between two noise time series was computed for all possible pairs of voxels (Fig 2.f.). Then, we averaged the Fisher’s z-transformed correlation coefficients [46] across all possible voxel pairs that include a single voxel of interest as follows:

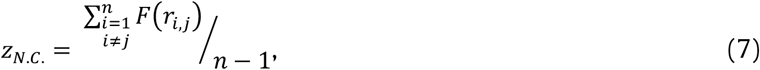

where *F*(*r*) = *tanh*^−1^(*r*) and *r_i,j_* is a Pearson’s correlation coefficient in noise time series between voxel *i* and *j*.

### Statistical analysis

We evaluated the statistical significance of tDCS effects, as follows: (1) the measures obtained in the pre-stimulation phase were subtracted from those obtained in the peri-stimulation and post-stimulation phases to control for tDCS-irrelevant variability in BOLD activity across daily sessions; (2) we evaluated whether (1) differ from ‘the corresponding subtracted measures of the sham-tDCS daily session’ to control for tDCS-irrelevant fluctuations of BOLD responses over time within single daily sessions. Specifically, for each measure, this evaluation was carried out using two mixed-effect ANOVA models, one for a-tDCS and the other for c-tDCS, because the polarity effects might not be symmetric [3]. See the supplementary materials for how we carried out the two robustness tests.

### Time course analysis of tDCS effects on the baseline, time-to-peak, and spatial-tuning measures of BOLD responses

#### Measurement

As for the baseline, we averaged the raw BOLD responses within each scan for each voxel. As for the time-to-peak, we fitted the sine function to the cycle-averaged BOLD time series. Then the phase value of the sine function was converted into the time unit (2*π*: 36 *sec*). As for the pRF width and pRF SI, we fitted the DoG function to the pair of traveling-wave scan runs, one traveling inward and the other outward.

#### Linear regression analysis

We did a linear regression analysis to test whether tDCS effects gradually increase or decrease during the peri-stimulation phase. We set 0 to regressors for pre-stimulation runs, [0.1, 0.3, 0.5, 0.7, 0.9] for peri-stimulation runs, and then 1 for post-stimulation runs so that we can interpret the slope of the regression as a degree of linear change during the stimulation.

### Voxel selection for Illustrative summary

We selected voxels with four criteria based on only the a-tDCS session data. First, the baseline parameter increased in the post-stimulation phase compared with pre-stimulation. Second, time-to-peak increased in peri-stimulation compared with pre-stimulation. Third, pRF width decreased in post-stimulation compared with pre-stimulation. Fourth, pRF SI increased in post-stimulation compared with pre-stimulation. 11% of voxels passed such criteria.

## Results

### Initial statistical results

During the peri-stimulation phase, a-tDCS affected only the temporal dynamics of BOLD responses to the transient visual input, decreasing the ‘peak amplitude’ (Z = −4.60, FDR-adjusted p < 1.E-05) and increasing the ‘time-to-peak’ (Z = 4.28, FDR-adjusted p < 1.E-04) and the ‘FWHM’ (Z = 3.34, FDR-adjusted p < 1.E-02) of the BOLD time series (Fig.3a). By contrast, no measure was affected by c-tDCS (Fig.3b).

**Fig. 3.**
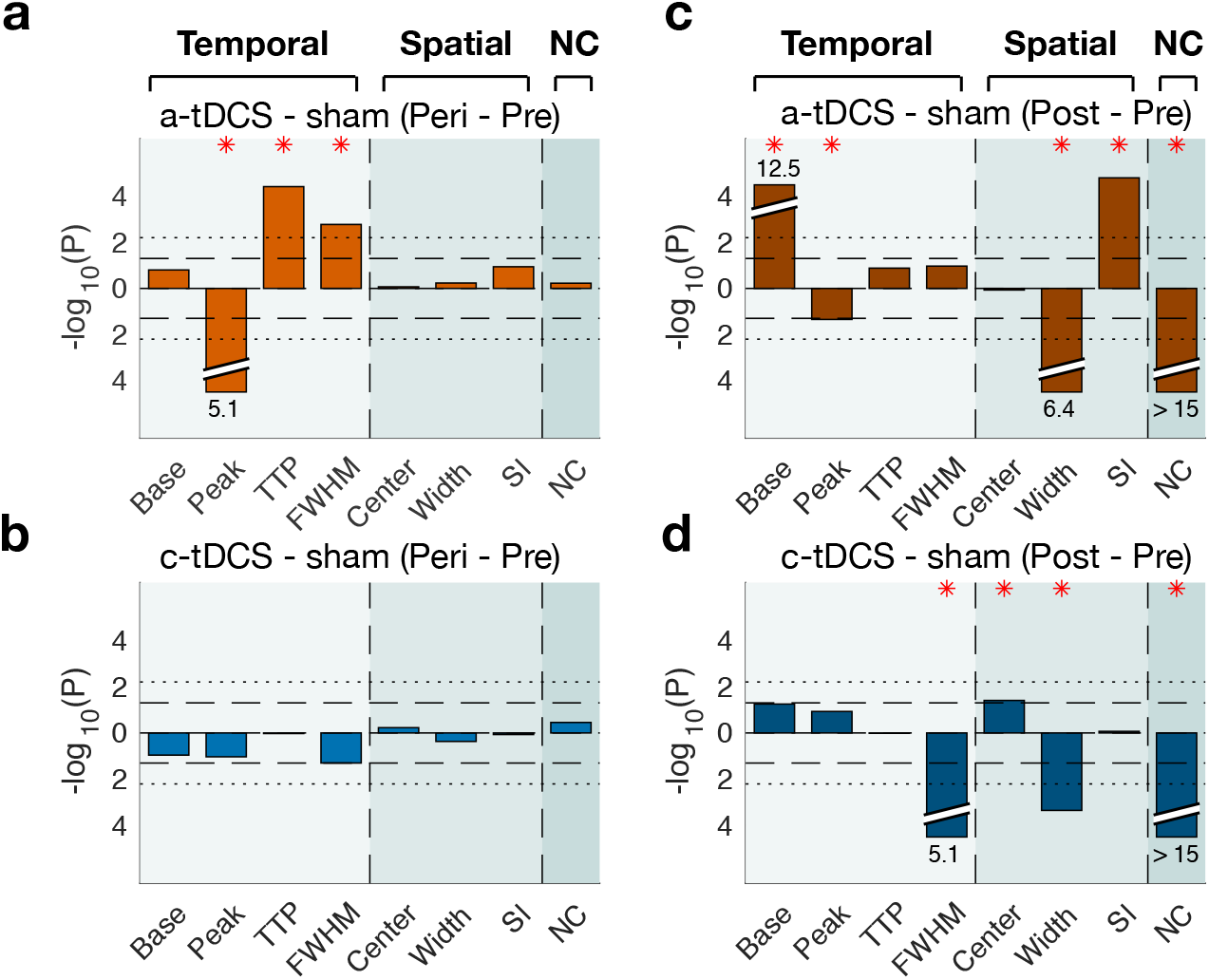
Initial statistical results (a-d) Modified Manhattan plots of mixed-effect ANOVA tests for the a-tDCS/peri-stimulation (a), c-tDCS/peri-stimulation (b), a-tDCS/post-stimulation (c), and c-tDCS/post-stimulation (d) effects. The signed +, increase; -, decrease) log of P values of the mixed-effect ANOVA tests are plotted against the temporal-profile (light teal background), spatial-profile (teal background), and noise-correlation (dark teal background) measures. Horizontal dashed and dotted lines demarcate the significance level with and without Bonferroni correction, respectively. Red stars mark the measures with significant (FDR-adjusted p < 0.05) tDCS effects.

During the post-stimulation phase, many of the BOLD measures were significant. Specifically, a-tDCS increased the baseline (Z = 7.38, FDR-adjusted p < 1.E-11), decreased the ‘peak amplitude’ (Z = −2.01, FDR-adjusted p < 0.05), reduced the pRF width (Z = −5.19, FDR-adjusted p < 0.01), increased the pRF SI (Z = 4.47, FDR-adjusted p < 0.01), and decreased the noise-correlation (Z = −12.90, FDR-adjusted p < 0.05) (Fig.3c). On the other hand, c-tDCS decreased the ‘FWHM’ (Z = −4.60, FDR-adjusted p < 0.05), increased the ‘pRF center’ (Z = 2.33, FDR-adjusted p < 0.05), decreased the ‘pRF width’ (Z = − 5.19, FDR-adjusted p < 0.05), and decreased the noise-correlation measure (Z = −12.90 FDR-adjusted p < 0.01) (Fig.3d).

Table 1 shows the detailed results of all (32 tests = 8 measures x 2 polarity x 2 stimulation phases) mixed-effect ANOVA tests.

**Table 1.**
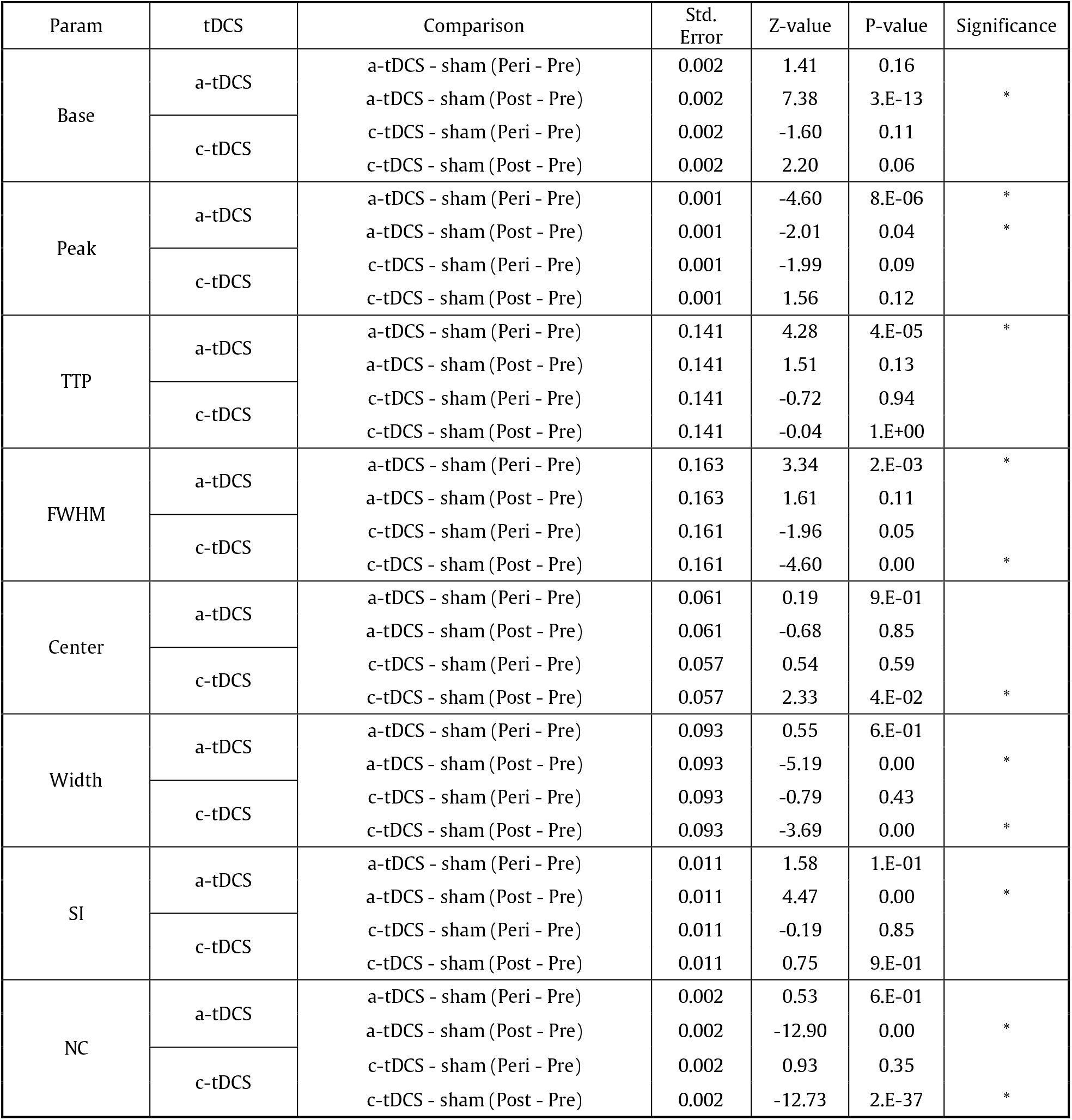
Initial statistical results from the planned comparison with Mixed ANOVA

### Evaluating the robustness of the initial statistical results

We assessed the robustness of the above significant effects by inspecting how reliably they retain statistical significance across different subpopulations of voxels or subjects. Such assessment is crucial in tDCS experiments on human subjects because the tDCS effects on human brains can be variable across local sites and subjects, although our tDCS stimulation protocols were optimized to the head anatomy of each subject.

First, we considered an imperfect alignment between daily sessions as the source of the across-voxel variability in tDCS effects. Using the between-session correlation in the BOLD time series during the pre-stimulation phase as a proxy of voxel correspondence, we refined the pool of valid voxels with varying levels of voxel correspondence. Then, we repeated the mixed-effect ANOVA test on those refined pools. Three of the initially significant results failed to survive this test (red symbols in Fig. 4c,d).

**Fig. 4.**
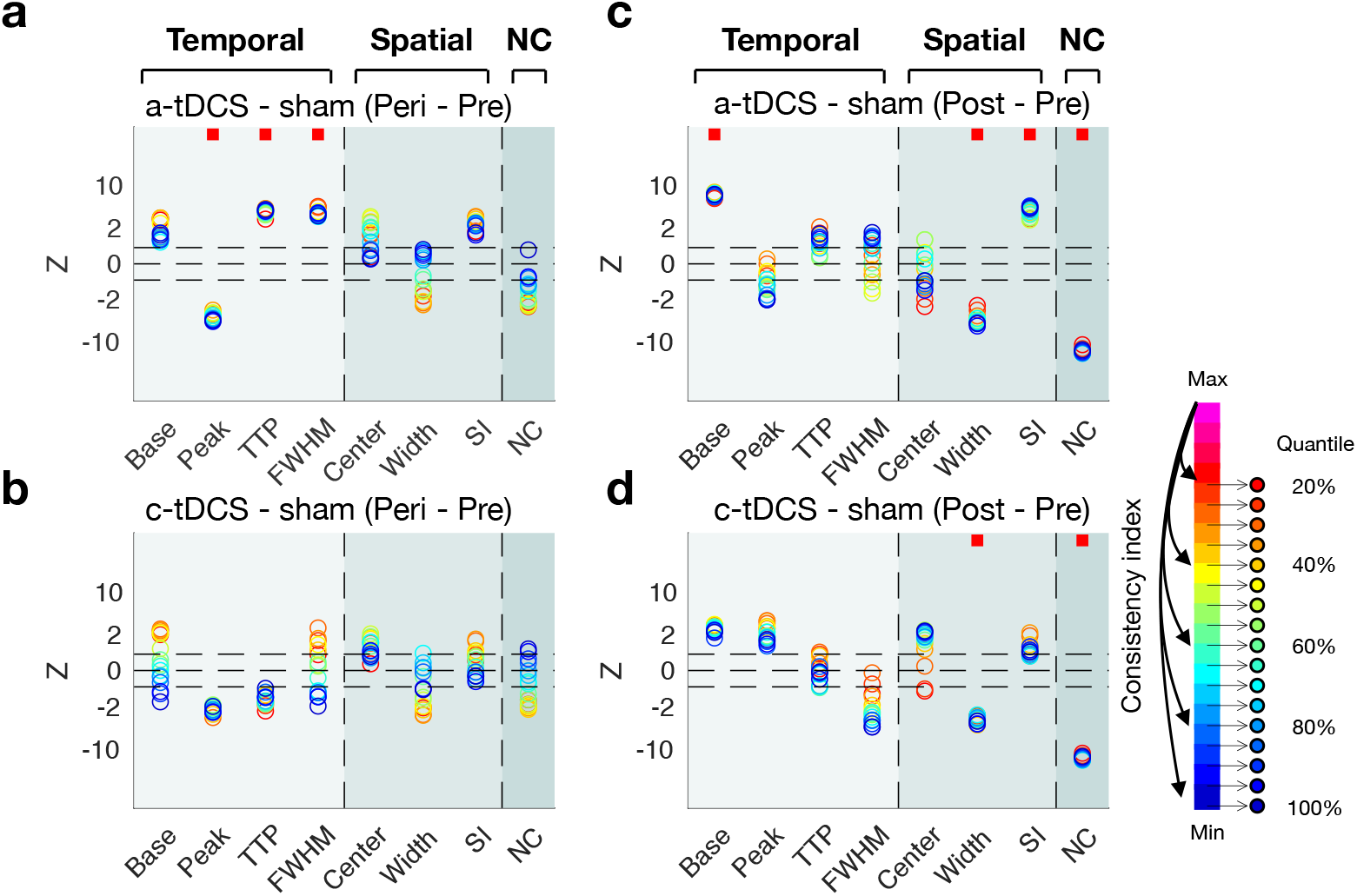
Test results of robustness to voxel misalignment (a-d) Plots of robustness-to-voxel-misalignment test results for the a-tDCS/peri-stimulation (a), c-tDCS/peri-stimulation (b), a-tDCS/post-stimulation (c), and c-tDCS/post-stimulation (d) effects. The test statics (Z; +, increase; -, decrease) are plotted against the temporal-profile (light teal background), spatial-profile (teal background), and noise-correlation (dark teal background) measures. Colors represent the quantiles of voxel-selection criteria for consistency index (equally spaced 17 bins from highest 20% to 100% with a step size of 5%). Horizontal dashed lines demarcate the significance level. Red squares indicate that the test statistics were significant at all bins.

Next, to assess the robustness of the statistical test outcomes to the subject-wise variability, we inspected whether the data set from a single ‘influential’ subject determined the statistical results using the jackknife resampling technique (Fig. 5). Seven of the initially significant outcomes failed to survive this test of robustness to subject-wise variability (red symbols in Fig. 5c,d).

**Fig. 5.**
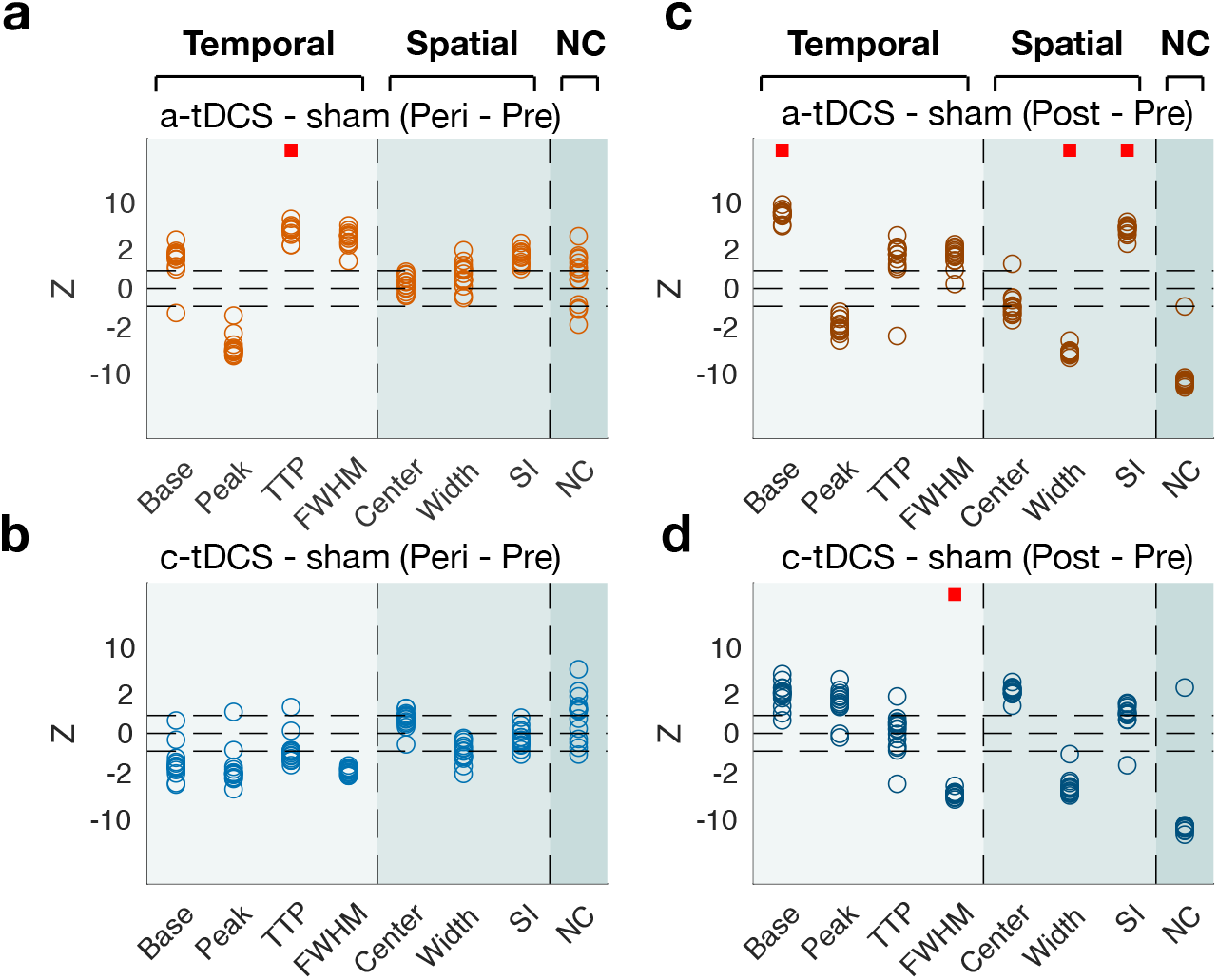
Test results of robustness to inter-subject variability (a-d) Plots of robustness-to-inter-subject-variability test results for the a-tDCS/peri-stimulation (a), c-tDCS/peri-stimulation (b), a-tDCS/post-stimulation (c), and c-tDCS/post-stimulation (d) effects. The statics (Z; +, increase; -, decrease) of the 15, n-1 jackknife resampling tests are plotted against the temporal-profile (light teal background), spatial-profile (teal background), and noise-correlation (dark teal background) measures. Horizontal dashed lines demarcate the significance level. Red squares indicate that the test statistics were significant throughout all tests.

By putting the initially significant tDCS effects to the two robustness tests, we found that only the anodal application of tDCS “reliably” modulated (i) the ‘time-to-peak’ measure of BOLD responses to the transient stimuli during the peri-stimulation phase, (ii) the ‘baseline’ measure of BOLD responses to the transient stimuli during the post-stimulation phase, and (iii) the ‘pRF width’ and ‘pRF SI’ measures of BOLD responses to the traveling-wave stimuli during the post-stimulation phase. In what follows, to see how these effects temporally develop, we probed the BOLD responses on a run-to-run basis.

### Time course of tDCS effects on BOLD baseline, time-to-peak, and pRF shape

For this run-to-run inspection, we acquired the baseline and time-to-peak measurements using the methods that can be applied not only to the BOLD responses to the transient visual input but also to those to the traveling-wave input (see Methods for details). We found that the baseline measures steadily grew while a-tDCS was applied and remained high after switching off the stimulation (red symbols and lines in Fig. 6.a). We did not find such a gradual elevation in the c-tDCS (blue symbols and lines in Fig. 6.a) or the Sham-control conditions (gray symbols and lines in Fig. 6.a).

**Fig. 6.**
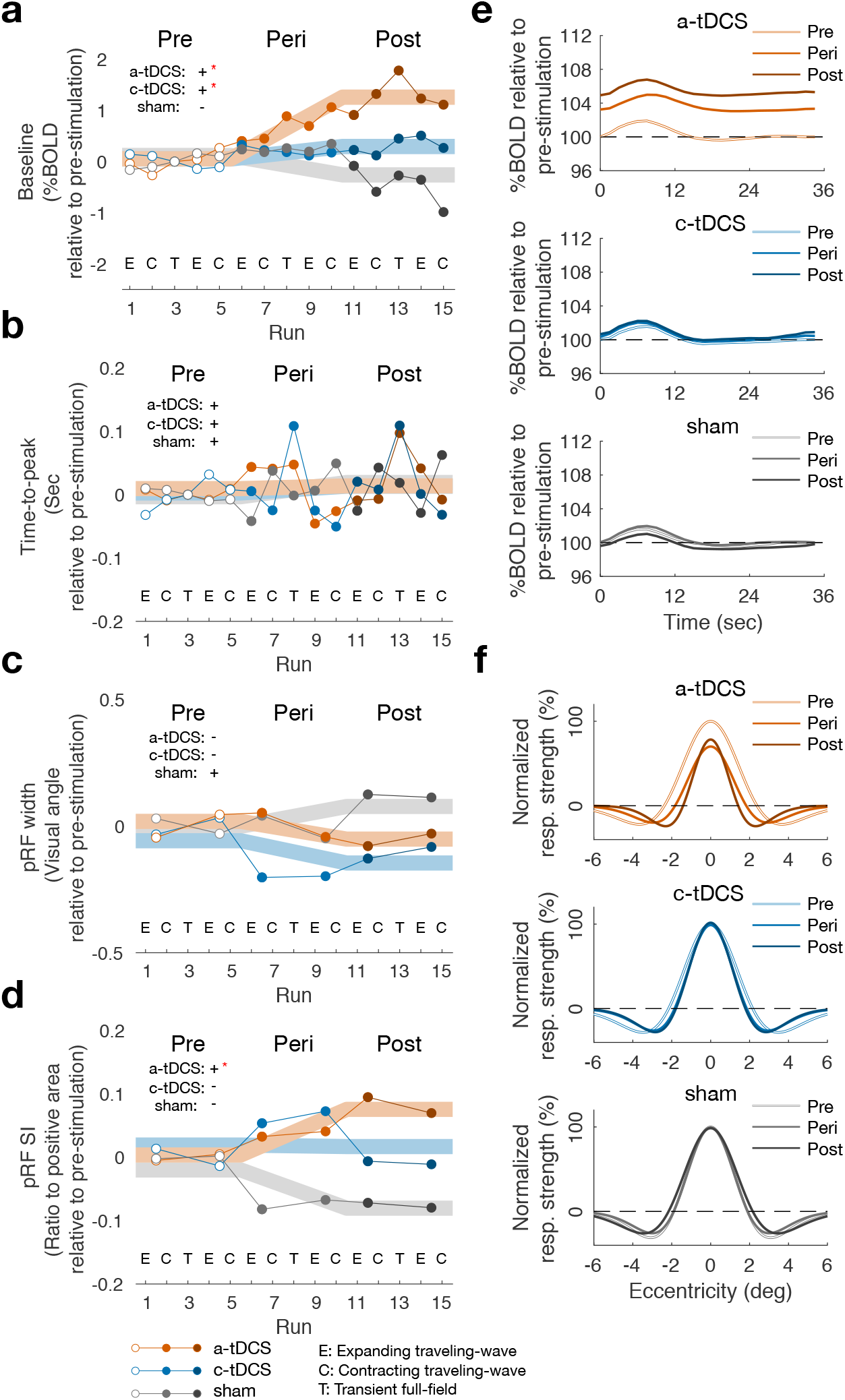
Time courses and illustrative summary of the principal tDCS effects (a-d) Across-run changes of baseline (a), time-to-peak (b), pRF width (c), and pRF SI (d) measures for a-tDCS (red), c-tDCS (blue), and sham-tDCS (grey) sessions. Across-voxel averages of the measures are plotted as a function of scan runs for each daily session. The measures were adapted to their own averages during the pre-stimulation phase. Dots and thin lines are the time courses of the data, whereas thick lines are those of the regression models, the signs and significances of which are indicated in the upper left corner of each panel (*: p < 0.05). Letters at the bottom denote the scan type: E and C for expanding and contracting traveling-wave input scans, respectively; T for transient whole-field input scans. (e-f) Illustrative summary of tDCS effects on BOLD responses to the transient full-field input (e) and pRF shapes (f). The plots summarize the results from a subset of voxels that showed the effects consistent with the grand average effects for all the following measures during the post-stimulation phase: ‘baseline’, ‘time-to-peak’, ‘pRF width’, and ‘pRF SI’. Panels correspond to daily sessions, indicated by the corresponding colors of plotted curves.

As for the time-to-peak measures, we fitted the sine function to the BOLD time series of each scan run by capitalizing on the fact that the transient and traveling-wave inputs share the same periodic cycle (see Methods for details). We did not find any trend in these time-to-peak measures— neither increase nor decrease—regardless of the tDCS stimulation conditions (Fig. 6.a).

In principle, the spatial-tuning (i.e., pRF) measures cannot be estimated from the BOLD responses to the transient visual input. In addition, the reliable pRF estimation requires the pair of traveling-wave scan runs, one traveling inward and the other outward. Thus, we acquired the spatial-tuning measures by fitting the pRF model to the BOLD time series in the two contiguous traveling-wave runs, which resulted in the six points of pRF width and pRF SI measures. The pRF-width and pRF-SI measures tended to decrease and increase, respectively, while a-tDCS was applied and remained high and low, respectively, after a-tDCS was withdrawn (red symbols and lines in Fig. 6.c,d).

In sum, from the moment of a-tDCS application, the BOLD responses steadily increased in baseline while their spatial-tuning function became narrow and suppressed in the surround.

### Illustrative summary of tDCS effects

Having identified the significant and robust effects of tDCS on visual BOLD responses and analyzed their time courses, we conclude the Results section by providing an *illustrative* summary of those effects. This illustrative summary is needed because it is not easy to appreciate the effects due to their modulatory (subtle), idiosyncratic (variable across individuals), and noisy (variable across voxels) nature. For this illustrative purpose, we opted to “cherry-pick” a specific set of voxels (see Methods for details). With this “paragon” set of voxels, we illustrate (1) the tDCS effects on the baseline and time-to-peak measures by plotting their cycle-averaged BOLD responses to the transient visual input (Fig. 6e) and (2) the tDCS effects on the pRF width and SI measures by plotting the pRF profiles with the averaged-across-voxel parameters best-fit to the BOLD responses to the traveling-wave input (Fig. 6f).

From (1), we can readily appreciate the impact of a-tDCS on the raw cortical responses to the transient input: it shifts their entire time series upward while delaying its peak slightly. From (2), we understand how a-tDCS changes the spatial profile of pRF: it sharpens the spatial tuning by narrowing the width of the positive center while augmenting the suppression at the immediate surround. These plots must be taken not as representative (typical) results but as a selected sample of data for an illustrative purpose.

## Discussion

To check how tDCS modulates cortical visual responses, we kept track of the spatial and temporal aspects of the BOLD activity of EVC before, during, and after tDCS application while varying the presence and polarity of tDCS application. We found that a-tDCS elevated the baseline BOLD activity without affecting the evoked activity while sharpening the voxels’ spatial tuning profiles by augmenting surround suppression. These effects grew gradually from electrical stimulation onset till its offset and remained substantial afterward. We ascertained the robustness of these findings to the two well-known nuisance factors of electrical brain stimulation studies.

### Buildup and persistence of a-tDCS effect on baseline BOLD activity

The current work found that the baseline BOLD activity gradually ramped up as the surface-positive current flowed into there and persisted for an extended period (at least 20 min) even after the current was switched off. Its magnitude was substantial, corresponding to 50% of the evoked BOLD activity (Fig. 6.a). This ramping and persisting dynamics are strikingly similar to that in the seminal work by Bindman and her colleagues [1,47], where the authors showed that when the polarizing current was applied for a 5 min or longer period, the spontaneous firing rate of somatosensory neurons increased steadily as the current was applied and persisted at that elevated level after it was switched off.

To our knowledge, our study provides the first report on the impact of a-tDCS on the baseline BOLD activity in human brains. Given its substantial magnitude and persistence in our study, one might wonder why such an impact has never been discovered in previous tDCS-fMRI studies. We note that most tDCS-fMRI studies focused on measuring the stimulus- or task-evoked, which led them to adopt the widespread practice of normalizing raw BOLD time series within each single scan run. This practice precludes the opportunity to observe the gradual changes in baseline activity. We anticipate that our finding on the baseline BOLD will likely be replicated by previous or future tDCS-fMRI studies if raw BOLD time series are normalized over the entire fMRI session as we did.

### Dissociation between baseline and evoked BOLD responses in tDCS effects on EVC

In previous studies, the surface-positive current modulated the evoked neural responses in the non-visual cortices in animals and humans, including the somatosensory cortex of rats [1,47], the motor cortex of cats [2], the Hippocampal slices of rats [48], and the motor cortex of humans [5,7]. By contrast, in our study, the amplitude of the visually evoked BOLD responses did not show any hint of the ramping or persistence dynamics. The slight delay in time-to-peak was the only a-tDCS-induced evoked effect that was both significant and robust. But this delay was relatively small in size (0.1 ~ 0.5 sec) and limited in time (significant only in the transient input scan run during the peri-stimulation phase), which is far from the magnitude and dynamics of the a-tDCS’s impact observed on the baseline BOLD activity.

In line with our study, a similar dissociation between spontaneous and evoked responses had been noted by one of the earliest studies, where the effects of the surface-positive current on the spontaneous and evoked response were directly compared between the motor and visual cortices of the same cats [2]. Although the current-induced modulation of the evoked responses was much weaker in the visual cortex than in the motor cortex, that of the spontaneous responses was equally pronounced in the two cortices. Almost two decades ago, one of the seminal tDCS studies on humans reported the tDCS-induced changes in the visual evoked potentials (VEPs) from the human visual cortex, measured with EEG on the skull [49]. However, a close inspection of this work indicates that the VEPs were modulated not positively by a-tDCS but negatively by c-tDCS, confirmed by a recent study [50]. In a similar vein, the same group [51] also reported that the applications of a-tDCS and c-tDCS to the human visual cortex facilitated and suppressed, respectively, the occurrences of TMS-induced visual phosphenes, which had been cited as the evidence for the elevated excitability of the visual cortex by a-tDCS. However, this report has been invalidated by recent studies that more rigorously evaluated the effects of a-tDCS for a larger number of subjects [52,53].

Our findings support an emerging view that stresses the fundamental differences in tDCS effects between the sensory cortex and the motor or associated cortices by demonstrating that the anodal application of tDCS to the visual cortex failed to elevate the amplitude of the evoked cortical activity but substantially increased the baseline activity.

### A-tDCS sharpens the spatial tuning of pRF by augmenting surround suppression

Visual neurons are known to be suppressed by the visual input presented in the surround of its receptive field, a phenomenon called ‘surround suppression’ ([54–57]; see [58]). The surround suppression is explained by the divisive normalization model, in which the surround exerts a divisive, inhibitory influence on the responses to the center [54,59]. The pRF-estimation models based on the DoG function or its variants have incorporated those configurations in mapping the spatial tuning functions of single fMRI voxels [45,60]. The pRF models fit to BOLD responses tightly reflected the center-surround configurations measured by the multi-unit activity [61].

Based on the above rationale, the a-tDCS-induced increase of SI in our study suggests that a-tDCS augments the surround suppression in the visual cortex, which consequently sharpens the spatial tuning of visual neurons. Thus, our findings offer an account of the a-tDCS-induced improvement in human visual acuity, a phenomenon considered counterintuitive [62]. According to our interpretation, the augmented inhibitory influence from the suppressive surround narrows the spatial tuning curve of visual neurons, conferring the visual system with precise sensory encoding [63].

### Possible neural mechanisms for tDCS effects on EVC

The most potent yet intriguing effects of a-tDCS in our study are the simultaneous buildup and persistence of the baseline—but not evoked—BOLD activity and the surround suppression. What neural mechanism(s) mediates the co-existence of these seemingly counteracting effects? Why does a-tDCS not elevate evoked BOLD responses in the early visual cortex, unlike what has been found in the motor cortex? We address these questions by relating the hierarchical gradients in excitatory and inhibitory recurrent connections [64] to the neural metabolism underlying BOLD responses [65]. In brief, we conjecture that a-tDCS to the primate visual cortex preferentially increases synaptic inhibition by upregulating the activity of a subtype(s) of GABAergic cells in the recurrent connections, which increases the local brain metabolism leading to an increase in net BOLD activity.

Recent large-scale investigations on the biophysical properties of primate brains, including humans, revealed that the recurrent connections systematically vary along the hierarchy of cortical areas in two critical aspects. First, the number of spines on pyramidal dendrites—a proxy of synaptic excitation strength—steeply increases along the hierarchy, being lowest in the early visual cortex, intermediate in the motor cortex, and highest in the prefrontal cortex [66]. Second, the relative proportions of subtype GABAergic interneurons vary along the hierarchy: those inhibiting the synaptic input to excitatory neurons are most dominant in the early visual cortex, whereas those inhibiting excitatory neurons’ output become increasingly dominant along the hierarchy [67].

These “hierarchical gradients” [64] imply that a-tDCS is likely to cause different outcomes in recurrent neural activity depending on the cortical site of stimulation. Given the substantial differences in those gradients between the visual cortex and the motor cortex, we argue that our findings in the visual cortex should not be considered conflicting with the previous findings in the motor cortex. The effects of a-tDCS in our study might have to do with the extreme position of the early visual cortex on the gradients of recurrent connections. It is likely that a-tDCS in the early visual cortex preferentially up-regulates the contribution of the synaptic-inhibition-controlling type of GABAergic neurons to the E/I balanced network. Then, this can account for the a-tDCS-induced augmentation of surround suppression. Next, unlike the output-controlling type of GABAergic neurons, the synaptic-inhibition-controlling type of GABAergic neurons consume metabolic energy substantially so that its energy consumption overcomes the decrease of metabolic consumption due to spiking reduction [68,69]. Then, this can account for the a-tDCS-induced increase in baseline BOLD activity.

The scenario presented above is only a speculative and *ad hoc* account. Nonetheless, it is critical to consider the substantial differences in E/I balanced network composition across different cortical areas when interpreting any given effects of tDCS. Such considerations will make tDCS an effective tool for investigating the circuit-level neural mechanisms responsible for major psychiatric diseases such as Schizophrenia. For instance, the previously reported weak surround suppression in schizophrenic patients [70,71] might be enhanced by a-tDCS on their visual cortex, which can be an exciting future extension of the current work.

