## Supplementary material for "Transcranial direct current stimulation elevates the baseline activity while sharpening the spatial tuning of the human visual cortex"

**Methodological details**

**Subjects exclusion**

While twenty-seven subjects initially participated in the current study, eight of them could not be used for analysis for the following reasons. Eight subjects had to quit in the middle of data collection due to schedule conflicts, which is understandable given the extended (more than two months) period of data collection (see “Experimental design” section for the details). Four subjects completed the entire participation but were excluded from data analysis because their head motion exceeded the spatial extension of a single voxel (2mm x 2mm x 2mm) in any of the fMRI scans conducted over four daily sessions.

**Double-blind procedure**

To blind experimenters to the type of a given tDCS session, we instructed experimenters to simply run a pre-designated code, which was labeled only for the date to be run.

**Display**

Visual stimuli were generated with MATLAB using the MGL on a Macintosh Mac Pro [1] and presented using an LCD projector Canon XEED SX60 (1400 × 1050 pixel resolution, 60 refresh rate) onto a projection screen inside the magnet bore (39.5 cm × 29.5 cm screen size, 98cm distance from the eyes, field of view 22° × 17° visual angle). Subjects viewed the projection screen via a mirror with a multilayer dielectric reflective coating installed on the head coil. We linearized the gamma function of the display using a luminance meter Konika Minolta LS-100.

**Fixation task**

Throughout all functional scan runs, we required subjects to perform a fixation task to fix their gaze at the fixation dot (0.05° in diameter) centered on the display and to control their attention at a constant level. They had to press a button whenever a pair of small green dots (0.05° diameter), which were constantly rotating along a red annulus (0.05° width, 0.175° radius) around the fixation dot, reversed the rotating direction.

**MRI acquisition**

MRI data were acquired using a Siemens MAGNETOM Trio, A Tim System 3T eco. At the beginning of each daily session, T1-weighted (MPRAGE) anatomical volume images were acquired (20 channel, 1.6 sec repetition time, 2.36 ms time to echo, 9 deg flip angle, 1 × 1 × 1 mm^3^ voxel size, 256 × 256 matrix size). In the retinotopy mapping session, high resolution T1 images were additionally collected (32 channel, 2.4 sec repetition time, 3.42 ms time to echo, 8 deg flip angle, 0.85 × 0.85 × 0.85 mm^3^ voxel size, 320 × 320 matrix size). Functional scan images were acquired using an Echo-plannar Imaging protocol: seven subjects were imaged with a single-band EPI protocol (20 channel, 1.5 sec repetition time, 30 ms time to echo, 75 deg flip angle, 2 × 2 × 2 mm^3^ voxel size, 96 × 96 matrix size, 24 slices); eight subjects were imaged with a multi-band EPI protocol (4 MB factor, 20 channel, .75 sec repetition time, 33 ms time to echo, 55 deg flip angle, 2 × 2 × 2 mm^3^ voxel size, 96 × 96 matrix size, 36 slices).

**Retinotopy mapping for ROI definition**

We defined the visual areas by running ten retinotopy mapping scans using the phase-encoding method [2]. A wedge-shaped dartboard pattern (3 cycles per visual angle degree(v.a.d.)) traversed the visual space by rotating around the fixation dot (10°/s) either clockwise (odd-numbered scan runs) or counter-clockwise (even-numbered scan runs). From the time series of BOLD responses to the rotating wedge, we estimated the temporal phase values for individual voxels and projected them onto the flattened surface segmented from T1 images using Freesurfer [3,4] (<http://surfer.nmr.mgh.harvard.edu/>). Based on the phase map, the early visual areas including V1, V2 and V3 were manually defined [5].

**fMRI data preprocessing**

The images acquired across different sessions and runs were aligned to one another in the following procedures. For each subject, we aligned the in-plane T1 images, which were acquired in each scan run, to the high resolution T1 images, which were acquired in the retinotopy mapping scan by finding the transformation matrices resulting in the highest quality alignment. Next, we aligned the functional images to the high resolution T1 images using the transformation matrices defined in the above procedure. The entire alignment procedures were done via SPM8 (http://fil.ion.ucl.ac.uk/spm) and mrTools (http://cns.nyu.edu/heegerlab/?page = software).

Correction for motion artifacts were done in the following procedures. The motion artifacts occurring within daily sessions were corrected using a motion compensation procedure [6,7]. To achieve the fine-grain match in spatial position between the voxels imaged in different days, we resampled high resolution T1 images with the functional voxel size (3 mm) and then aligned the functional voxels from each daily session to the nearest resampled voxels only when the distance between two were within voxel size.

We detrended the functional scan time series for the low-frequency non-physiological component [8] by subtracting the time series convolved with one cycle duration (36 sec) boxcar function [9]. When defining the hemodynamic impulse response function (HIRF), we used the cycle-averaged time series in raw BOLD scale. The differences in acquisition time between slices were corrected.

**Voxel selection**

The voxels included in data analysis were determined in the following steps. Since our tDCS protocol focuses on the early visual cortex as target areas, data analysis was confined to the voxels within the ROIs of V1, V2, and V3, which were defined by the retinotopy mapping scans (see “Retinotopy mapping for ROI definition” for details). Next, we screened out the voxels whose BOLD responses are likely to be contaminated by the non-neuronal, physiological noises such as those associated with the magnetic field inhomogeneity induced by respiration and cardiac pulsation. Specifically, based on the tight linkage between temporal fluctuations of BOLD responses and the fraction of variance explained by the physiological noises[10,11], we computed the temporal standard deviation (tSTD) of the residual BOLD responses after regressing the detrended BOLD time series (see “fMRI data preprocessing” for details) onto the cycle-averaged BOLD responses for each voxel and discarded the voxels whose tSTD values fell on the highest 2% of the entire ROIs for a given subject. Next, we further screened out the voxels (i) when the goodness of fit (r2) of a model was <0.1 for either the temporal profile scan runs or the spatial profile scan runs, or (ii) when the SNR was lower than the criterion. The SNR was defined as the amplitude of stimulus presentation frequency component divided by the average amplitude of frequency components three times higher than the stimulus presentation frequency in pRF mapping scan. The SNR criterion was determined as the s95th percentile of SNR value from the noise ROI (white matter, cerebrospinal fluid) voxels for each subject.

**Electric field simulation to optimize tDCS for individuals**

*Individual head models*. Sophisticated 3D finite element (FE) head models were constructed from T1-weighted MR images using a series of software packages including SimNIBS, FreeSurfer, and FSL [12–14]. The FE head models were composed of five tissue compartments including white matter, gray matter, cerebrospinal fluid, skull, and scalp, and their electrical conductivities were assumed to be 0.126, 0.276, 1.65, 0.01, and 0.465 S/m, respectively [15]. Interfacial boundaries of the head models were visually inspected and manually corrected, if needed, using ITK-SNAP [16]. The numbers of nodes and elements of the constructed FE models are presented in Table 1. We then modelled eight circular sponge electrodes and attached them on the scalp surface of the head models. The electrodes were attached at Oz, PO7, PO8, PO3, PO4, Pz, O9, and O10, according to the extended 10-20 EEG placement system [17]. The electric conductivity of the electrodes was assumed to be 1 S/m, and their radius and thickness were set to 10 mm and 5 mm, respectively.

*Analysis of electric field generated by multichannel tDCS*. The electric potential V generated by tDCS is governed by the following Laplace equation:

∇∙(σ∇V) = 0,	(1)

where $\sigma$ is the electrical conductivity [18]. In our study, we adopted finite element method (FEM) with first-order tetrahedral elements to numerically solve (1). Then, the current density J can be readily found using

J = σ E = σ (-∇V),	(2)

where E is electric field intensity. Electric field intensity inside the head due to multi-channel tDCS can be evaluated as the linear sum of electric field intensity values due to all possible electrode pairs. When the first electrode (Oz) was defined as the reference electrode, the total electric field intensity $E_{total}$ can be evaluated as

$E_{total}= \sum_{n=2}^{8} I_{n}E_{n}$,	(3)

where $I_{n}$ is the electric current flowing between the reference and n-th electrodes, and $E_{n}$ is the electric field intensity due to the unit current between the reference and n-th electrodes [19,20]. To obtain the electric potential distribution, two different Dirichlet boundary conditions such as 0 V and 1 V were applied on the reference and the n-th electrodes, respectively. The electric field intensity was then scaled by integrating the electric current density passing through the electrode using (2), resulting in $E_{n}$.

For the optimal stimulation of V1, we used evolutionary strategy (ES) to minimize the objective function defined as

I = arg min $\sum_{k=1}^{N} {(\left\| E_{total}^{k} \right\|^{2}-e_{target}^{k})}^{2}$	,	(4)

where

$e_{target}^{k}=\left\{ \begin{aligned} 1, if k-th node \in V1 \\ 0, elsewhere \end{aligned} \right.$	(5)

and $E_{total}^{k}$ is the electric field intensity at the k-th node. Programs for FEM and ES were coded with Fortran90 under Microsoft Visual Studio 2015 environment, compiled by Intel Parallel studio 2017 Composer Edition, and executed in Windows10 installed on Intel i7-7700 personal computer system.

Supplementary Table 1. The number of nodes and elements in extracted models.

| Subjects | Head volume mesh | | Cortex surface mesh | |
| --- | --- | --- | --- | --- |
|  | # nodes | # elements | # nodes | # elements |
| 1 | 292190 | 1726439 | 106171 | 212338 |
| 2 | 275265 | 1623743 | 72420 | 144836 |
| 3 | 277385 | 1646402 | 71789 | 143574 |
| 4 | 304713 | 1824272 | 83059 | 166114 |
| 5 | 280943 | 1663645 | 70150 | 140296 |
| 6 | 387018 | 2275979 | 115585 | 231166 |
| 7 | 299751 | 1786332 | 70131 | 140258 |
| 8 | 650590 | 3642552 | 122588 | 245172 |
| 9 | 708395 | 4149576 | 175862 | 352504 |
| 10 | 670983 | 3764211 | 125202 | 250400 |
| 11 | 637819 | 3572342 | 119213 | 238422 |
| 12 | 684744 | 3847269 | 123922 | 247840 |
| 13 | 662161 | 3718189 | 125355 | 250706 |
| 14 | 655035 | 3671579 | 122573 | 245142 |
| 15 | 656032 | 3677142 | 128865 | 257726 |

**tDCS protocol**

We applied tDCS using the Starstim tES 8-channel system with a multi-channel MRI extension kit. The system was remotely controlled by the software Neuroelectrics Instrument Controller (NIC2). Eight saline-soaked circular sponges (8 cm^2^) with carbon rubber electrodes were arranged to form a center-surround geometry on the surface of the head (Fig 1.d). The total amount of electric current was set to 2mA, with the amounts of current for individual electrode being optimized respectively for individual subjects based on the results of the electric field simulation. The major fraction of electric current (> 1.5mA) was applied through the main electrode that was located at the center. The electrode arrangement and current amount were identical for a-tDCS and c-tDCS: only the direction of current was reversed between the two stimulation conditions. The durations of ramping-up and ramping-down electric current were set to 10 seconds each (Fig 1.c). In all types of stimulation phases, we started to ramp up current as fMRI imaging got started. We started to ramp down current as soon as ramping-up was done in the shame stimulation phases but after the fMRI scan run was completed in the genuine stimulation phases. Each subject received a total of 18-minute long (5 run × 216 sec) genuine tDCS in both a-tDCS and c-tDCS sessions.

**Electric field prediction for BOLD data analysis**

We manually identified the actual positions of the electrodes in the in-plane T1 images for each daily session. Based on those positions, we predicted the electric filed (EF) intensity for each daily session using the same FEM used above. Then the EF intensity value of each node was projected onto voxels. To estimate the EF intensity of the voxels located deep below the surface, we fitted the three-dimensional Gaussian function to the across-subject average raw EF intensity:

$EF estimate=a+ G\left( \left( x,y,z \right), \Sigma\right), \Sigma= \left( \begin{matrix} C_{xx} & C_{xy} & C_{xz} \\ C_{xy} & C_{yy} & C_{yz} \\ C_{xz} & C_{yz} & C_{zz} \end{matrix} \right)$,	(9)

where $a$ is a constant, $\left( x,y,z \right)$ is a relative coordinate from the main electrode

and $\Sigma$ is a covariance matrix. Then we obtained EF estimate for individual voxels based on the fitted 3D Gaussian function and their relative coordinates from the position of the main electrode (Fig 1.e).

**Temporal-profile analysis of BOLD responses to transient visual stimuli**

*Stimulus*. To see whether the application of tDCS incurs any change in the temporal dynamics of BOLD activity in the early visual cortex, we repeatedly presented brief visual stimuli and inspected diverse aspects of the temporal profile of BOLD responses to those stimuli. In each 36-second long cycle of the transient-stimulus scan runs, sinusoidal radial gratings were briefly presented, only for 3 seconds, within a large (8° in radius) circular aperture then disappeared for a prolonged period of time (33 sec) (Fig 2.a.). The spatial frequency of radial gratings was optimized for the given eccentricity range [21], changing logarithmically over the eccentricity such that it was about 3 cycle per v.a.d. at the fovea and 1 cycle per v.a.d. at the eccentricity of 8 v.a.d. The orientation and spatial frequency of the gratings were matched to those used in traveling-waves-stimulus scan runs.

*Model fitting*. We modeled the hemodynamic impulse response function (HIRF) with the two gamma function [22]. Then we fit the HIRF to the time series of BOLD responses of each voxel by predicting the time series based on the convolution of the stimulus event matrix with the HIRF. The best-fitting parameters of the HIRF were identified as those resulting in the maximum correlation between the predicted and observed time series of cycle-averaged BOLD responses. Then the scale parameters were analytically solved using the least square method, which is mathematically equivalent to minimizing the sum of squared error (SSE) but more efficient and less vulnerable to the local minima problem [23].

**Spatial-profile analysis of BOLD responses to travelling-wave visual stimuli**

*Stimulus*. To see whether the application of tDCS incurs any changes in spatial tuning property of the early visual cortex, we presented a ring-shape visual stimulus and let it expand or contract slowly over the visual field and inspected diverse aspects of the spatial profile of BOLD responses to those stimuli. In each 36-second long cycle of the travelling-wave-stimuli scan runs, sinusoidal radial gratings that were presented within ring-shape apertures slowly (4.5 v.a.d. per second) traversed the visual field in the radial direction (Fig. 2.b.). The spatial frequency of radial gratings was optimized for the given eccentricity range [21], changing logarithmically over the eccentricity such that it was about 3 cycle per v.a.d. at the fovea and 1 cycle per v.a.d. at the eccentricity of 8 v.a.d. The ring-shape aperture had the width of 2/3 v.a.d. and moved smoothly with the velocity 8 v.a.d. per 36 sec, so any spatial point was filled with stimuli 3 seconds, which was matched to that of the transient-stimuli scan run.

*Model fitting*. We modeled the pRFs of single voxels with the one-dimensional (1D) difference-of-Gaussian (DoG) function [24] because we drifted the traveling-wave stimuli only along the eccentricity dimension and wanted to capture the potential tDCS effects not just of a facilitatory nature but also of an inhibitory nature. With the DoG function, pRF is defined as a gain function ($g(x)$) of stimulus input ($x$) falling on a 1D space (eccentricity in our case) as follows:

$g\left( x \right)= {\beta_{+} g}_{+}\left( x \right)- {\beta_{-} g}_{-}\left( x \right)$;

$g_{+}\left( x \right)=\exp-\left( \frac{\left( x-x_{0} \right)^{2}}{2{\sigma_{+}}^{2}} \right)$; $g_{-}\left( x \right)=\exp-\left( \frac{\left( x-x_{0} \right)^{2}}{2{\sigma_{-}}^{2}} \right)$	,	(6)

where $g_{+}\left( x \right)$ and $g_{-}\left( x \right)$ are positive and negative, respectively, Gaussian functions, the respective contributions of which to $g\left( x \right)$ are expressed by the two coefficients, $\beta_{+}$ and $\beta_{-}$; $x_{0}$ is the center of pRF, and $\sigma_{+}$ and $\sigma_{-}$are the standard deviations of the positive and negative Gaussian functions, respectively. Next, we predicted the aggregated responses of a population of visual neurons within a voxel to the traveling-wave stimuli ($r(t)$) by convolving a binary matrix of spatio-temporal stimulus input ($s\left( x,t \right)$) with $g\left( x \right)$ (Fig 2.c-e.). Finally, we predicted the BOLD time series by convolving $r(t)$ with the HIRF defined by the transient-stimulus scan. Importantly, the HIRF used in this convolution was defined separately for individual voxels by the transient-stimulus scan in the same stimulation phase in which the traveling-wave-stimulus scans of interest were run. In this way, any differences in pRF shape for a given voxel—its spatial tuning property—between the different tDCS stimulation phases cannot be ascribed to possible changes in its temporal profile—its temporal dynamics property—between the corresponding tDCS stimulation phases. As was done for HIRF fitting, the best-fitting parameters of pRF were those resulting in the maximum correlation between the cycle-averaged predicted and observed BOLD time series. Likewise, the scale parameters were analytically solved by the least square method, which ensures that the pRF parameters are independent of the temporal-profile measures of ‘baseline’ and ‘peak amplitude’.

**Statistical analysis**

In sum, based on the fMRI data analyses described above, we obtained a total of eight different measures—four temporal-profile measures, three spatial profile measures, and one noise correlation measure (Fig. 2)— for each of the nine stimulation phases—3 tDCS daily sessions x 3 stimulation phases (Fig. 1.a). In evaluating the statistical significance of tDCS effects on these eight measures, we exploited the sham-controlled crossover design in the following steps. First, to control for any tDCS-irrelevant variability in BOLD activity across daily sessions, for each daily session data set, we subtracted the measures obtained in the pre-stimulation phase from those obtained in the peri-stimulation and post-stimulation phases. Next, to control for any tDCS-irrelevant fluctuations of BOLD responses over time within single daily sessions, we evaluated whether or not ‘the “subtracted” measures of the peri-stimulation and post-stimulation phases of the a-tDCS and c-tDCS daily sessions’ differ from ‘the corresponding “subtracted” measures of the peri-stimulation and post-stimulation phases of the sham-tDCS daily session’. Specifically, for each of the eight measures, this evaluation was carried out using two mixed-effect ANOVA models, one for a-tDCS and the other for c-tDCS, because the polarity effects might not be symmetric[25], and thus the anodal and cathodal conditions are better to be treated rather separately.

*Mixed-effect ANOVA models*. In each model, one fixed effect, the ‘tDCS-versus-sham’ contrast, is evaluated with the two nested random effects, the ‘voxel’ variable nested in the ‘subject’ variable. We opted to build such a hierarchical mixed-effect model because the tDCS effects are likely to vary across voxels and subjects owing to variation in miscellaneous factors including EF strength, partial gray-matter volume, head anatomy, age, and attention level, which makes individual voxels and subjects that nest those voxels the nuisance variables whose potential interferences with the fixed effect should be taken into account. We did a planned comparison because our purpose is to test whether the tDCS effects are significant or not rather than to figure out what factors are significant. We implemented the mixed-effect ANOVA models in R (version 3.6.2), while controlling the false discovery rate (FDR) at an $\alpha$ level of 0.05 using the Benjamini-Hochberg procedure [26]. As for the test statistics, we opted for Z-test, rather than T-test, because the sample variance could be regarded as the population variance due to the sufficiently large sample size (i.e., the number of individual voxels nested in subjects).

*Evaluating the robustness of the ANOVA results to voxel selection criteria based on consistency*. As described earlier (see ‘Voxel selection’ section), the pool of voxels included in analysis was determined using a very conservative set of selection criteria. However, we considered a possibility that the procedure of aligning functional images across different daily sessions is imperfect and thus subject to a certain degree of error. If that is the case, the observed differences in the measures between the tDCS and sham conditions might not be entirely ascribed to the genuine tDCS effects. To address this issue, we came up with a new criterion for voxel selection based on the consistency in BOLD responses during the pre-stimulation phase between the daily sessions. The rationale behind this criterion is as follows: given that the impact of imperfect alignment varies across voxels depending on their locations, the extent to which voxels corresponded to one another between daily sessions can be assessed by measuring how similar their BOLD time series during the pre-stimulation phase were between daily sessions. For individual voxels, we indexed the consistency in BOLD responses during the pre-stimulation phase between the daily sessions by averaging the Pearson’s correlation between the cycle-averaged BOLD time series from the corresponding functional scan runs. Based on those consistency indices, we varied the voxel-selection criterion from highest 20% to 100% in increments of 5%, resulting in a total of 17 different pools of voxels. Then, we repeated the mixed-effect ANOVA test for those 17 pools of voxels to check whether the statistical results were invariant to those voxel selection criteria.

*Jackknife resampling*. To check the possibility that the voxels sampled from an outlying subject disproportionately contribute to the observed statistical outcomes, we created $N_{subject}$ (the number of subjects used for data analysis, which is 15) datasets from the original dataset by successively omitting the voxels from each subject at a time using the jackknife resampling technique. We repeated the mixed-effect ANOVA test on those 15 jackknifed datasets.

**Time course analysis of tDCS effects on the baseline, time-to-peak, and spatial-tuning measures of BOLD responses**

*Measurement*. As for the baseline, we averaged the raw BOLD responses within each scan (120 frames) for each voxel. The scale was normalized as a percent-change relative to the same value from same kind of runs in pre-stimulation phase of same daily session. To compare the net change regardless of stimuli kind, we subtracted the averaged value from same kind of runs in pre-stimulation phase. As for the time-to-peak, we fitted the sine function to the cycle-averaged BOLD time series. The best-fitting phase parameter was identified as those resulting in the maximum correlation between the predicted and observed time series of BOLD responses. Then the phase value of sine function was converted into the time unit($2\pi:36 sec$). Then, we subtracted the averaged value from same kind of runs in pre-stimulation phase. As for the pRF width and pRF SI, we fitted DoG function to the pair of traveling-wave scan runs, one traveling inward and the other outward. There were two pairs for the each stimulation-phase. The fitting procedure was same with the section ‘Spatial-profile analysis of BOLD responses to travelling-wave visual stimuli’. We subtracted the averaged value from the two pairs of pre-stimulation phase. For each parameter, we averaged the values from voxels to obtain one data point for one scan run.

*Linear regression analysis*. To test whether tDCS gradually increase or decrease parameters during peri-stimulation phase, we did simple linear regression analysis. We set 0 to regressors for five pre-stimulation runs, [0.1, 0.3, 0.5, 0.7, 0.9] for five peri-stimulation runs, and then 1 for five post-stimulation runs so that we can interpret the slope of the regression as degree of linear change during the stimulation. For each parameter, we did regression with voxel-averaged value so there were fifteen data point for baseline and time-to-peak parameter and six data point for pRF width and SI parameter for each tDCS session. The p-value of significance test was 0.05.

**Illustrative summary**

*Time series of BOLD response to the transient visual input*. The cycle-averaged BOLD response to the transient visual input was normalized as a percent-change relative to the baseline measure in the pre-stimulation phase, so that the baseline measures of the peri-stimulation and post-stimulation phases could be fairly compared between the three daily tDCS sessions. Then we averaged the BOLD time series from voxels for each stimulation phase.

*pRF profiles with the averaged parameter*. We averaged four pRF model parameters ($\beta_{+},\beta_{-},\sigma_{+},\sigma_{-}$) that captures the width and SI of pRF from selected voxels. The $x_{0}$ parameter was fixed to 0. The peak response amplitude was normalized as a percent relative to the same measure in the pre-stimulation phase, so that the response amplitude of the peri-stimulation and post-stimulation phases could be fairly compared between the three daily tDCS sessions.
